# HCN channels reveal conserved and divergent physiology in supragranular pyramidal neurons in primate species

**DOI:** 10.1101/2025.08.22.671856

**Authors:** Cristina Radaelli, Matthew Schmitz, Xiao-Ping Liu, Scott Sawchuk, Ximena Opitz-Araya, Mark Hudson, Naz Taskin, Darren Bertagnolli, Jeff Goldy, Andrew L Ko, Benjamin L Grannan, Jason S Hauptman, Anoop P Patel, Charles Cobbs, Kimberly A Smith, William J Spain, Ed S Lein, Trygve Bakken, Nikolai C Dembrow, Jonathan T Ting, Brian E Kalmbach

## Abstract

The physiological properties of human and rodent neurons differ, yet the extent to which these differences reflect human specializations is often unclear. Compared with their rodent counterparts, human supragranular pyramidal neurons possess enriched HCN-channel-dependent intrinsic membrane properties and a related sensitivity to synaptic inputs containing delta/theta band frequencies. We tested whether other primate species possess enriched HCN-channel dependent membrane properties. We found ubiquitous *HCN1* subunit gene expression in supragranular glutamatergic neurons across New World Monkeys, Old-World Monkeys, and great apes in single nucleus RNA-sequencing datasets. Using Patch-seq recordings from acute and cultured brain slices, we found robust HCN-dependent physiological properties in supragranular pyramidal neurons in a species of New-World monkey (*Saimiri sciureus*) and two species of Old-World Monkey (*Macaca mulatta, Macaca nemestrina*). In both human and macaque neocortex, HCN-related intrinsic properties increased in magnitude with increasing laminar depth, especially in one transcriptomic cell type. Within this type, HCN dependent properties were more pronounced in macaque than human neurons. These findings indicate that HCN-governed membrane properties and sensitivity to delta/theta band frequencies are roughly conserved in supragranular pyramidal neurons across at least 36 million years of primate evolution.

## Introduction

Although gross neocortical cytoarchitecture is roughly conserved among terrestrial mammals, the expansion of the supragranular layers (layers 1-3) in humans, marks a divergence from many species, including commonly studied rodent species ^1–5^. Additionally, the supragranular layers of human neocortex have more diverse cell sizes and many unique cellular properties compared to their rodent counterparts ^23^. High throughput single cell/nuclei RNA-sequencing (sc/sn RNA-seq) has facilitated the development of comprehensive cell type taxonomies that permit matched comparisons of the cell type composition of human versus mouse neocortex ^6–9^. These analyses have revealed greater cell type diversity overall in human neocortex, including cell types found in human which are not present in mouse supragranular layers, and greater depth from pia dependent variation in gene expression among glutamatergic neurons ^10^.

Although these differences may reflect human specializations, they could also represent more general features of primate neocortex.

In addition to differences in cytoarchitecture, cell number and type composition, human supragranular pyramidal neurons possess several distinct biophysical and synaptic properties in comparison to their rodent counterparts. Human supragranular pyramidal neurons have more complex dendrites than rodent neurons in terms of total dendritic length, number of branches and spine density. These features enable a greater number of input synapses and enhanced information processing ^11,12^. Additionally, human dendrites may have unique spikes that enable computations not found in rodent dendrites ^13^. Local, recurrent synaptic connectivity is also unique in the human supragranular layers, where synaptic connections between human neurons are larger and more reliable^14–17^. Human supragranular pyramidal neurons also exhibit a directed acyclic network architecture, facilitating information flow in a hierarchical manner, in contrast to rodent neurons ^18^.

Ion channel expression differences between human and murine neurons have also been consistently reported to contribute to physiological distinctions between species ^17,19,20^. For example, hyperpolarization-activated cyclic nucleotide-gated ion channel (HCN channel) gene expression and associated intrinsic membrane properties are enriched in human supragranular pyramidal neurons compared with their mouse counterparts ^12,21–23^. HCN channels conduct an inward current that slowly activates upon hyperpolarization and slowly deactivates upon depolarization ^24,25^. Consequently, HCN channels counteract changes in membrane potential and together with passive properties of the cell membrane, promote subthreshold membrane resonance and the integration of synaptic input containing delta and theta frequency bands ^22,26–29^. HCN channels also counteract capacitive filtering, ensuring that the time course of synaptic responses measured at the soma are similar regardless of their site of origin on the dendrite ^26,30^. In the human neocortex HCN-dependent membrane properties increase in magnitude with increasing laminar depth in a single transcriptomic cell type, a feature that is not observed in rodent supragranular neurons ^10,14,22,31^.

However, it remains unclear whether enriched HCN properties reflect unique human adaptations or whether they represent characteristics of primate supragranular pyramidal neurons in general. Here, we tested whether HCN conductance contributes to the intrinsic physiology of supragranular glutamatergic neurons in non-human primate species. We examined HCN channel gene expression in supragranular glutamatergic neurons of various primate species and performed Patch-seq ^32–36^ experiments in pig-tailed macaques, rhesus macaques and squirrel monkeys. Patch-seq experiments enabled us to directly compare HCN dependent properties of human and non-human primate neurons in homologous cell types.

## METHODS

### Non-human primate specimens

All procedures involving non-human primates were approved by the University of Washington’s Institutional Animal Care and Use Committee (IACUC) and conformed to the NIH’s Guide for the Care and Use of Laboratory Animals. Brain tissue was obtained from animals designated for the Washington National Primate Research Center’s Tissue Distribution Program. Tissue availability was limited to three non-human primate species (*Macaca nemestrina, Macaca mulatta, Saimiri sciureus*) and two cortical areas (Temporal Cortex, Motor Cortex). We obtained data from 67 pig-tailed macaques (n = 28 male, n = 39 female), 4 rhesus macaques (n = 2 male, n = 2 female), and 6 squirrel monkeys (n = 2 male, n =2 female, n= 2 unknown). The donors were aged 2-22 years old altogether (mean = 9.04 ± 0.09). There were no differences in HCN-dependent properties between male and female subjects (Effect of sex on sag ratio: p= 0.93; effect of sex on resonance frequency: p= 0.13; effect of sex on input resistance: p= 0.64, Independent samples t-test).

### Human specimens

We gained access to resected adult human brain tissue collected during neurosurgery via a collaborative network of four local hospitals (Swedish Medical Center, UW Medical Center, Harborview Medical Center, and Seattle Children’s Hospital). All patients provided informed consent, and experimental procedures were approved by the hospital Institutional Review Boards prior to the start of the study. We collected surgical tissue from the Middle Temporal Gyrus (MTG) localized distal to the pathological tissue from patients undergoing surgery for treatment refractory epilepsy or tumors.

This study includes unpublished data from 22 donors aged 18-72 years (mean = 41.6± 2.06), totaling 66 recordings (see Supplementary Table 1 for human donor metadata), as well as a reanalysis of previously published data from ^10^. The latter dataset comprises 61 patients aged 19 to 63 years (mean age: 37.1 ± 1.8 years), amounting to 385 recordings. There were no differences in HCN-dependent properties between male and female subjects (Effect of sex on sag ratio: p= 0.44; effect of sex on resonance frequency: p= 0.39; effect of sex on input resistance: p= 0.36, Independent samples t-test).

### Brain slice preparation

Non-human primates were anesthetized with isoflurane gas prior to transcardial perfusion of carbogenated (95% O2/5% CO2) artificial cerebrospinal fluid consisting of (in mM): 92 N-methyl-D-glucamine (NMDG), 2.5 KCl, 1.25 NaH_2_PO_4_, 30 NaHCO_3_, 20 4-(2-hydroxyethyl)-1-piperazineethanesulfonic acid (HEPES), 25 glucose, 2 thiourea, 5 Na-ascorbate, 3 Na-pyruvate, 0.5 CaCl_2_·4H_2_O and 10 MgSO_4_·7H_2_O; NMDG aCSF). After exsanguination, the cerebrum was extracted and neocortical areas of interest including Primary Motor (M1), Superior Temporal Gyrus (STG) and Inferior Temporal Gyrus (ITG) were dissected and placed in chilled, carbogenated NMDG aCSF for transport to the Allen Institute for further processing or in sucrose aCSF (in mM: 2.5 KCl, 1.25 NaH_2_PO_4_, 25 NaHCO_3_, 0.5 CaCl_2_, 7 MgCl_2_, 7 dextrose, 205 sucrose, 1.3 ascorbic acid, and 3 sodium pyruvate, bubbled with 95% O_2_/5% CO_2_ to maintain a pH of 7.4) for processing at the University of Washington.

Tissue blocks were oriented to maximize the preservation of the neocortical layers and the intactness of pyramidal cell dendrites. Tissue slabs were sectioned at 300-350 μm using NMDG aCSF or sucrose aCSF on a vibrating blade microtome (VT1200S, Leica Biosystems or Compresstome VF-300, Precisionary Instruments; Vibratome 3000). Brain slices were transferred to a carbogenated and warmed (34°C) chamber containing NMDG aCSF for 10 minutes, with the exception of tissue sections made in sucrose aCSF, which were transferred to a carbogenated warmed (34°C) holding aCSF (in mM: 125 NaCl, 2.5 KCl, 1.25 NaH_2_PO_4_, 25 NaHCO_3_, 2 CaCl_2_, 2 MgCl_2_, 10 dextrose, 1.3 ascorbic acid, and 3 sodium pyruvate) for 30 min and then left at room temperature until recording.

Slices cut in NMDG aCSF and intended for acute recorbdings, were stored in a carousel prefilled with room temperature aCSF consisting of: 92 mM NaCl, 2.5 mM KCl, 1.25mM NaH_2_PO_4_, 30mM NaHCO_3_, 20mM HEPES, 25mM glucose, 2mM thiourea, 5mM Na-ascorbate, 3mM Na-pyruvate, 2 mM CaCl_2_·4H_2_O and 2 mM MgSO_4_·7H_2_O. Slices reserved for tissue culture were transferred to membrane inserts (PICMORG, Millipore) in 6 well plates containing 1mL each of culture media (8.4 g/L MEM Eagle medium, 20% heat-inactivated horse serum, 30 mM HEPES, 13 mM D-glucose, 15 mM NaHCO_3_, 1mM ascorbic acid, 2mM MgSO_4_·7H_2_O, 1 mM CaCl_2_.4H_2_O, 0.5 mM GlutaMAX-I and 1 mg/L insulin). We adjusted the culture media to osmolality 300-310 mOsmoles/kg and to pH 7.2 to 7.3. We changed the media approximately every two days until slices were used for Patch-seq experiments. The cultured slices were stored in a humidified, 35°C, 5% CO_2_ incubator for at least a few hours and up to a maximum of 8 Days In Vitro (DIV). As such, cultured slices extended the life of tissue specimens and maximized tissue use. Cultured brain slices also enabled us to utilize adeno associated viral (AAV) genetic tools to label pyramidal neurons with a fluorescent reporter protein. In these cases, cultures were infected by direct application of concentrated AAV viral particles on the slice surface 1-12 hours after plating.

### Adeno-associated virus (AAV) vectors preparation, cloning and packaging

We used two AAV vectors previously demonstrated to label pyramidal neurons in macaque *ex vivo* brain slices - AiP11633 (pAAV-hsA2-AiE0140h-140h-minRho-SYFP2-WPRE3-BGHpA) and AiP12787 (pAAV-AiE0140h_3xC2minBG-SYFP2-WPRE3-BGHpA). The latter enhancer is an optimized 3x core concatemer of the enhancer element contained in AiP11633. These tools enabled us to record from labeled cells within 3-5 days post infection at the earliest. Full plasmid sequences and maps are available at Addgene (plasmid IDs: 163503 and 191726); ^37,38^.

Following a previously established protocol ^37,38^, we prepared the PHP.eB serotype packaging plasmid. We then transfected 15µg of the AAV plasmid together with 15 ug capsid plasmid (PHP.eB) and 30 ug pHelper plasmid (Cell Biolabs) into one 15-cm plate of confluent HEK-293T cells using PEI Max (Polysciences Inc., catalog # 24765-1), to generate crude AAV virus preparations.

We followed previously developed protocols ^37–39^ and made a few changes to the co-transfection mixture preparation. One day after transfection, we changed the medium to low serum (1% FBS). Three days post-infection we collected cells and supernatant, destined to be freeze-thawed three times, to release AAV particles. Afterwards, we applied benzonase nuclease (MilliporeSigma catalog # E8263-25KU) for 1 hour to degrade free DNA. Following centrifugation at 3000g for 10 minutes, we isolated the supernatant and concentrated it to approximately 150 µL by using an Amicon Ultra-15 centrifugal filter unit (NMWL 100 kDa, Sigma #Z740210-24EA) at 5000g for 30-60 min, yielding a titer of approximately 3-5.0E+13 vg/mL.

### Patch-seq recordings

Prior to Patch-seq experiments, we cleaned surfaces with RNAse Zap (Sigma-Aldrich), followed by nuclease-free water. We pulled glass patch clamp electrodes with an open tip resistance of 3 to 6 MΩ when filled with internal solution containing (in mM): 110.0 K-gluconate, 10.0 HEPES, 0.2 EGTA, 4 KCl, 0.3 Na_2_-GTP, 10 phosphocreatine disodium salt hydrate, 1 Mg-ATP, 20 mg/mL glycogen, 0.5U/mL RNase inhibitor (Takara, 2313A), 0.5% biocytin and 0.02 Alexa 594 or 488 – pH adjusted to 7.3 with KOH.

We performed whole cell patch-clamp recordings using either an Axoclamp 2B or a Multiclamp 700B amplifier and custom acquisition software written in Igor (MIES https://github.com/AllenInstitute/MIES or custom scripts written by Rick Gray/N. Dembrow).

Electrical signals were digitized at 50 kHz by an ITC-18 (HEKA Elektronik, Lambrecht/Pfalz, Germany) and were digitally filtered at 10 kHz. We compensated pipette capacitance and balanced the bridge (5-30 MΩ) during all current clamp recordings. All recordings were collected at a controlled temperature of 32-34°C. We provided continuous fresh ACSF perfusion at a rate of 4mL/min through the recording chamber that contained the following ingredients (in mM): 119 NaCl, 2.5 KCl, 1.25 NaH_2_PO_4_, 24 NaHCO_3_, 12.5 glucose, 2 CaCl_2_·4H_2_O and 2 MgSO_4_·7H_2_O (pH 7.3-7.4). In some of our recordings (n= 102), we suppressed fast synaptic transmission with either 25 µM d-APV, 20 µM DNQX and 5 µM GABAzine or 20 µM DNQX, 25 µM dAPV and 100 µM picrotoxin. We analyzed and compared the *ex vivo* recording conditions in macaque cells, and we did not find significant differences in intrinsic neuronal properties in cells recorded with or without blockers (effect of blockers on sag ratio: p= 0.63; effect of blockers on resonance frequency: p= 0.08; effect of blockers on input resistance: p= 0.51, Independent samples t-test).

In a subset of experiments in temporal cortex (n=19), we bath applied an HCN channel blocker, ZD7288 (10 µM; Tocris) for 10-12 minutes and compared HCN dependent properties before and after application. ZD7288 was prepared from frozen concentrated stock solutions and diluted in recording aCSF.

After completing the recording, we applied negative pressure (40-100 mbar) to aspirate the neuron’s nucleus into the patch pipette. We then expelled the cellular content into a tube containing a reverse transcription buffer, and stored the samples at –80°C. The nucleated Patch-seq technique offers enhanced transcriptomic data quality by improving RNA yield and minimizing degradation, as demonstrated in previous studies ^40^.

### Recording protocol

We applied two current injection protocols to measure HCN channel-dependent properties of supragranular pyramidal neurons. The first stimulus set consisted of a series of 1-s square wave current steps that increased from-200 pA to 50 pA in 50 pA step increments. We scaled this current protocol up or down to accommodate neurons with low or high input resistances, respectively. Maximum and steady-state input resistance were calculated from the linear portion of the current/voltage relationship elicited in response to the stimulus. Voltage sag was defined as the ratio of maximum to steady state input resistance. To exclude cases in which the maximum input resistance was lower than the steady state input resistance, sag ratio values < 0.95 were excluded from analysis because a steady-state voltage response did not occur during the 1s current step.

The second protocol was a chirp stimulus which was a constant amplitude sinusoidal current that increased in frequency logarithmically from 0.2-40Hz over 20 s. For each neuron, the chirp amplitude was adjusted to produce a peak-to-peak voltage deflection of ∼10 mV. The impedance amplitude profile (ZAP) was determined by taking the ratio of the Fourier transform of the voltage response to the Fourier transform of the chirp: *|z(f)|=√((Re(Z(f)))^2^+(Im(Z(f)))*^2^) where Im(Z(f)) and Re(Z(f)) are the imaginary and real parts of the impedance Z(f), respectively. The frequency at which the peak impedance was observed was referred to as the resonance frequency (*f*_R_), while the 3 dB cutoff was defined as the frequency at which the ZAP signal was attenuated to a value of *√(1/2)Zmax*. Electrophysiological features were calculated using a modified version of the IPFX Python package or using customs scripts written in Igor Pro.

### Processing of Patch-seq samples

We employed the SMART-Seq v4 Ultra Low Input RNA Kit for Sequencing developed by Takara (634894) to perform reverse transcription of RNA into cDNA throughout 20 PCR cycles.

Following cDNA amplification, libraries were constructed using Nextera XT DNA Library Preparation Kit (Illumina FC-131-1096). We sequenced samples to ∼500k-1 million paired-end 50b reads/sample. Non-human primate paired-end reads were aligned to the Mmul_10 rhesus macaque reference genome, while human paired-end reads were aligned to the GRCh38.p14 reference genome.

### Patch-seq mapping

To identify the corresponding transcriptomic cell type for each Patch-Seq sample, we utilized a correlation-based mapping approach. For each neuron, we calculated the correlation of its gene expression profile with a set of marker genes defining each branch of a consensus MTG cross-species taxonomy derived from several non-human primate species ^41^. For a correspondence of human Patch-seq samples mapped to this taxonomy versus the Hodge et al., 2019 MTG taxonomy see Fig. S1. Briefly, L2/3 IT_2 has a ∼1:1 correspondence to the Exc L3-5 RORB COL22A1 type. For the other types there is a many-to-many relationship, but the L2/3 IT_1 type seems to specifically capture a bigger fraction of the deep L3 (Deep Exc L2-3 LINC00507 FREM3 and Exc L3-4 RORB CARM1P1) types. We included Patch-seq samples in the dataset with ≥2000 genes detected, a correlation score ≥0.45, a normalized sum “on” type maker gene expression (nms) ≥0.3, and at least 20% of reads aligned to introns.

### Gene expression analysis

We accessed previously published MTG single nucleus single cell RNA-seq data from human and non-human primate neocortex ^41^. The mouse single cell RNA-seq cortical data, including somatosensory, visual, auditory and motor cortex, were instead extracted from ^42^.

Cell subclass identities were transferred from the cortical cross-species dataset to the mouse and spatial transcriptomic datasets using the Allen Institute cell type mapper (https://knowledge.brain-map.org/mapmycells/process/).

MTG spatial transcriptomics data was accessed from ^41^. Cortical depths were determined programmatically by manually annotating the inner and outer boundaries of the cortical plate. Then for each point, the nearest point on the opposing boundary was found to form a radial line. Cells were given a cortical depth from 0 to 1 corresponding to the point on the closest radial by perpendicular distance. Normalized *HCN1* values were then plotted against these cortical depths for each cell. Rolling expression means for each subclass were computed for cells within a window of 0.05 cortical depth.

## Statistical Analysis

Data wrangling and analysis were performed in IgorPro, Python or R. For depth from pia-dependent analysis, we fit HCN channel-dependent membrane properties with a multiple regression model followed by a two-way Analysis of Variance (ANOVA), and the calculated p-values were FDR adjusted for multiple comparisons using the Benjamini/Hochberg method. For other comparisons we assessed statistical significance through two-way ANOVA, t-tests and illustrated comparisons between groups using box plots and regression plots. For effect sizes we report partial eta-squared (η²ₚ) or R^2^. In each figure caption we reported the appropriate statistical analysis and results.

## RESULTS

### *Ubiquitous HCN1* expression in supragranular glutamatergic neurons in primate species

We first asked whether genes encoding HCN channel pore forming subunits (*HCN1-HCN4)* ^24^ and an auxiliary protein, TRIP8b^43^ (*PEX5L*), are expressed in supragranular glutamatergic neurons in several primate species. The *PEX5L* gene encodes TRIP8b, an auxiliary protein that co-localizes with HCN1, promoting HCN channel trafficking and surface expression, as well as modulating HCN gating, kinetics, and function^43^. We quantified HCN-related gene expression in glutamatergic neurons in existing snRNAs-seq datasets obtained from New World monkeys (marmoset), Old World monkeys (rhesus macaque), great apes (gorilla and chimpanzee), and humans (Fig 1a ^41^). To compare HCN channel-related gene expression to a rodent species we also quantified mouse gene expression from the Allen Brain Cell Atlas ^42^. In each primate species *HCN1* and *PEX5L* were abundantly expressed in many neurons in each glutamatergic subclass, whereas in mice expression of these genes was less pervasive (Fig 1a). The expression of genes encoding other subunits was relatively low in each subclass/species (Fig 1a ^41^). In each species we compared *HCN1* expression in supragranular glutamatergic neurons to expression in L5 extratelencephalic projecting neurons (L5 ET), which are known to be enriched for HCN-related membrane properties across various mammalian species ^19,22,44,45^. In mouse neocortex, *HCN1* expression was approximately two times higher in L5 ET neurons compared with supragranular glutamatergic neurons (Fig. 1b). In contrast, in each primate species *HCN1* expression was more similar in L5 ET neurons and supragranular glutamatergic neurons (Fig 1b). *HCN1* expression was also similar in each type of supragranular glutamatergic neuron in each primate species (Fig. S2). These observations suggest that there is extensive *HCN1* expression in supragranular glutamatergic neurons in several higher primate species as opposed to rodents. Importantly, the *HCN1* subunit, which displays the fastest activation kinetics among the four pore-forming subunits, is associated with band-pass filtering of delta/theta band frequencies^46^.

**Figure 1.**
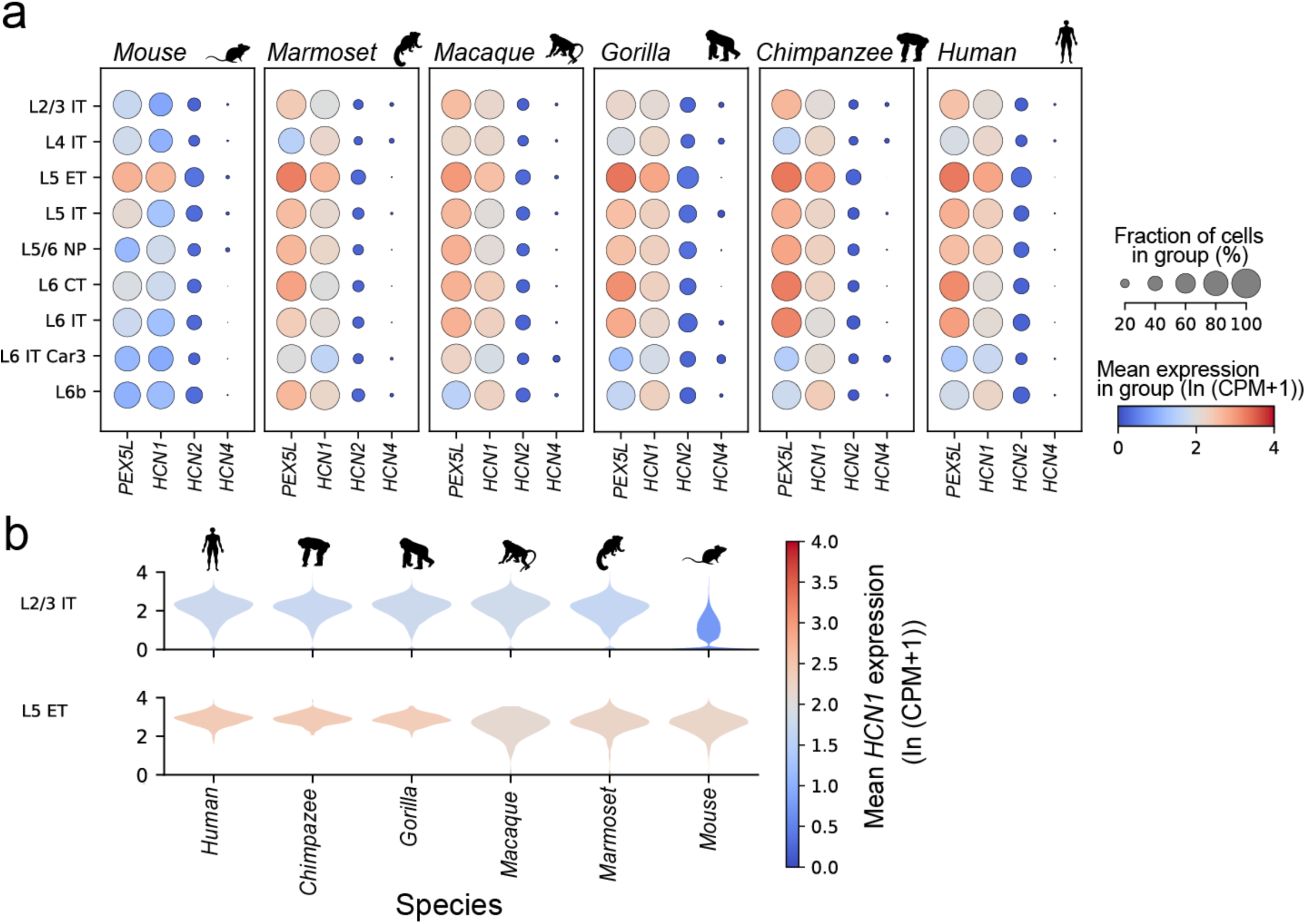
HCN channel-related gene expression is enriched in supragranular glutamatergic neurons in various primate species. a) Dot plots of *HCN1*, *HCN2*, *HCN4* and *PEX5L* expression in six species: mouse, marmoset, macaque, gorilla, chimpanzee and human. Colors and size of the dot plot correspond to the mean gene expression and the fraction of cells expressing the gene for the group, respectively. b) Violin plot of *HCN1* expression in layer 2/3 IT and L5 ET neurons for each species. Colors denote mean group expression.

### Supragranular pyramidal neurons in New World and Old Word monkeys possess HCN channel-dependent membrane properties

We next tested whether the HCN channel-related gene expression we observed in primate supragranular neurons is paralleled by HCN-dependent intrinsic membrane properties. To accomplish this, we obtained living brain tissue from three species of non-human primates (Fig. 2a,b), through a collaboration with the Tissue Distribution Program at the Washington National Primate Research Center. We obtained Patch-seq/patch clamp recordings (n = 583) from acute and cultured brain slices in temporal and motor areas in each species which permitted us to test whether supragranular pyramidal neurons had HCN-dependent properties in New World Monkeys and Old-World Monkeys, two groups which diverged from a common ancestor approximately 36-50 million years ago ^47–49^ (Fig. 2a).

**Figure 2.**
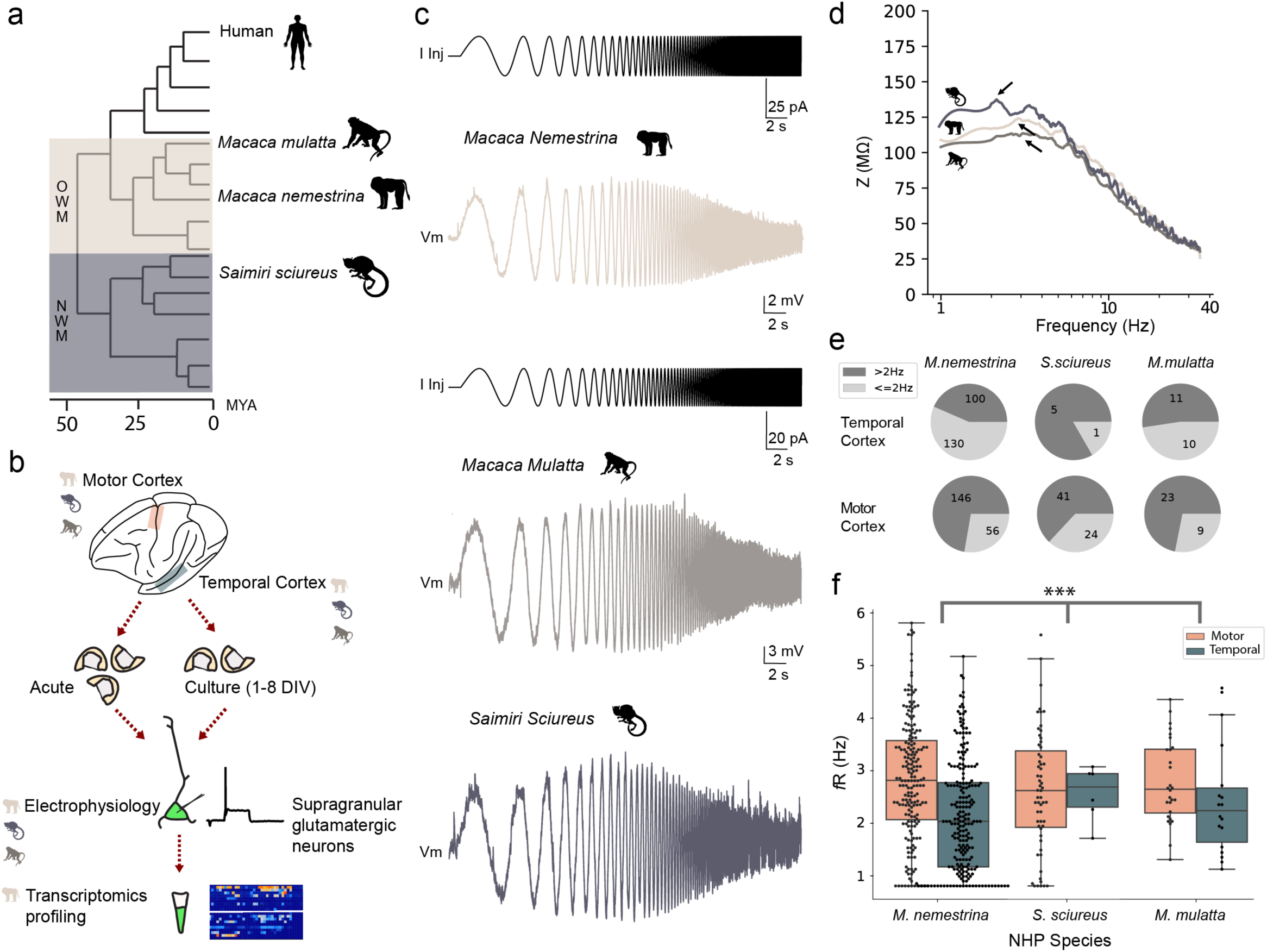
Supragranular pyramidal neurons in Old World and New World monkeys display subthreshold membrane resonance. a) Primate phylogenic tree showing the evolutionary distance among species, expressed in millions of years (MYA). b) Schematic of the experimental workflow. Tissue was resected from both temporal and motor neocortical areas for *ex vivo* slice preparation and patch-clamp recordings. From a subset of experiments, the nucleus was extracted for RNA sequencing. c) Example voltage response to a chirp current injection for each species. The same chirp stimulus applies to *M.mulatta and S.sciureus* examples. d) Impedance amplitude profile for these same cells. Arrows denote the resonance frequency for each neuron. e) Number of neurons in each species and neocortical area that had a resonance frequency greater than 2Hz. f) Resonance frequency of recorded cells across areas (temporal and motor) and species (*M.nemestrina*, *S.sciureus, M.mulatta*). Resonance frequency was higher in motor cortex for all species (denoted by **, p = 5.82e-07) and did not differ between brain slice preparation (p = 2.48e-01) or species (p = 6.25e-01); 3-Way ANOVA with species, preparation and area as the factors (Supplementary Table 2).

For each recording, we applied current injection protocols that detect the presence of HCN-conductance ^27,22,23,50^. One sign of HCN conductance is the presence of membrane resonance in the 2-14 Hz range in response to time varying input currents when measured near the typical resting membrane potential of neocortical neurons ^24,51^. To measure membrane resonance, we injected a chirp stimulus, which consisted of a constant amplitude sinusoidal current that increased in frequency logarithmically from 0.2 to 40 Hz over 20s (Fig. 2c). In each non-human primate species and neocortical area, we observed neurons with a resonance frequency (*f*_R_) greater than 2 Hz, but the proportion of cells exceeding 2 Hz tended to be higher in motor cortex (Fig. 2d,e). Resonance frequency was slightly higher in motor cortex neurons, and we did not observe significant differences between species (Fig. 2f, Fig. S3, Supplementary Table 2). A voltage “sag” in response to hyperpolarizing current injections is also indicative of the presence of HCN conductance ^24^. We observed neurons with clear sag in each species and in each neocortical area (Fig. S3). Finally, because HCN channels contribute to the resting conductance, we measured steady-state input resistance and 3db cutoff (a measurement of low pass filtering) for each neuron. Input resistance was similar across species and areas whereas 3db cutoff was higher in motor areas in all species (Fig. S3). These data suggest that HCN channel dependent properties are widespread in supragranular pyramidal neurons in primate species.

### HCN dependent features vary as a function of somatic depth from pia in macaque temporal cortex

In human middle temporal gyrus (MTG), HCN-dependent properties vary as a function of the somatic depth from the pial surface such that neurons in deep layer 3 have especially pronounced HCN dependent properties, compared to minimal HCN dependent properties observed in L2 pyramidal neurons ^10,22,23^. To test if this relationship holds true in macaque neocortex, we plotted several HCN-dependent properties as a function of the somatic depth from pia for each recording. We fit each feature with a multiple linear regression model in which depth from pia and slice preparation were the predicting variables. For this analysis we focused on data collected from the pig-tailed macaque where the sample size was largest (Fig. 3a).

**Figure 3.**
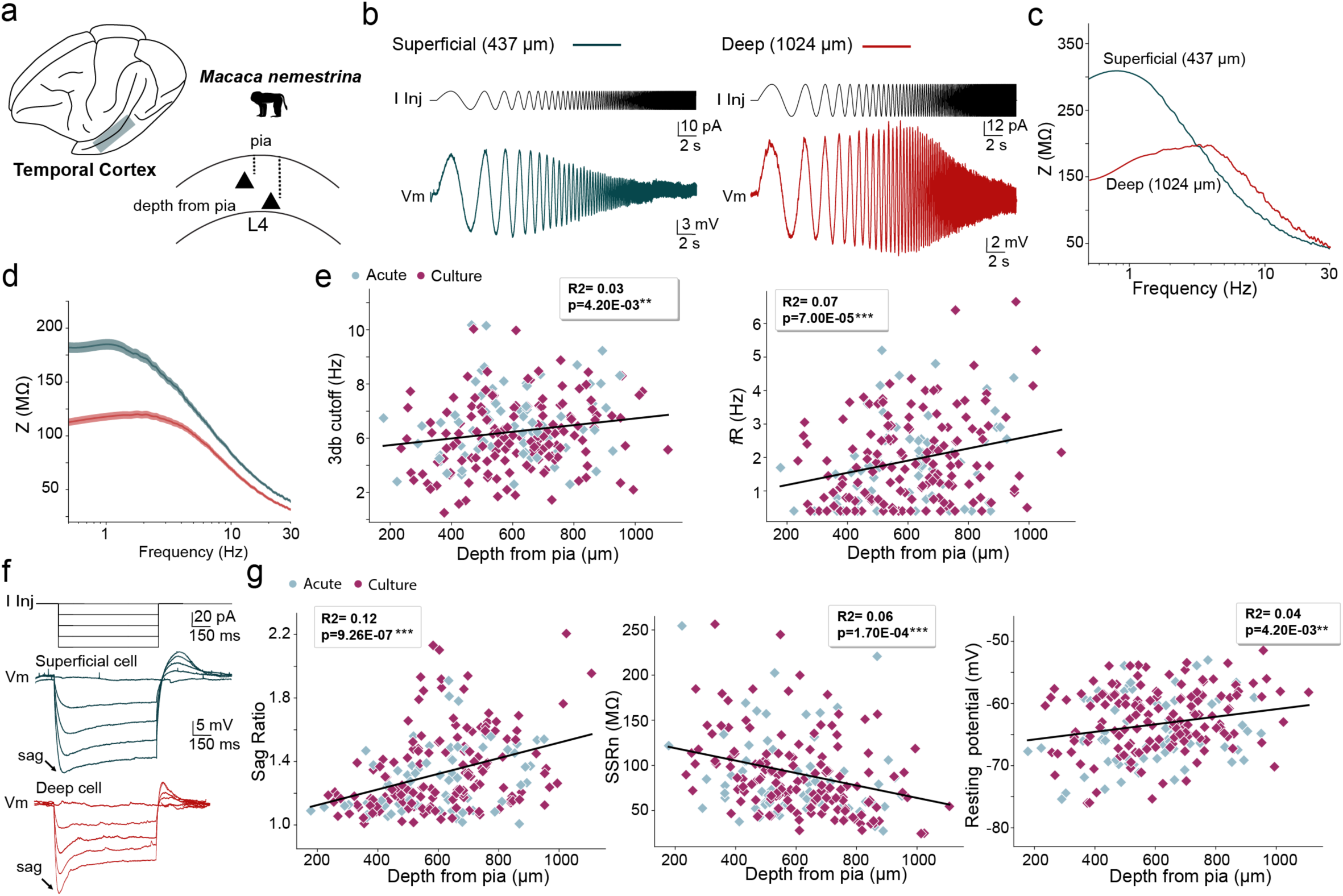
HCN-related intrinsic membrane properties are dependent on somatic depth from pia in the temporal cortex of the pig-tailed macaque. a) We quantified the subthreshold features of supragranular pyramidal neurons throughout the radial depth of the temporal cortex in *Macaca nemestrina*. b) example voltage response to a chirp stimulus for a superficial and deep neuron and c) corresponding impedance amplitude profiles. d) impedance amplitude profiles averaged across the superficial most one third (green) and deepest one third of neurons (red). e) 3dB cutoff and resonance frequency plotted as a function of distance from pial surface. f) Example membrane response to a series of hyperpolarizing current injections for a superficial and deep neuron. g) Sag ratio, steady-state input resistance and resting potential plotted as a function of distance from the pial surface. All p and R-squared values reflect the depth from pia effect from a 2-way ANOVA (*Effect on 3db cutoff: p=4.20e-03, on resonance frequency: p=7.00e-05, on SSRn: p= 1.70e-04, on resting membrane potential p= 4.20e-03, on sag ratio: p=9.26e-07;* Supplementary Table 3).

In the temporal cortex several HCN-related membrane properties varied as a function of the somatic depth from the pial surface (Fig. 3; Supplementary Table 3). Resonance frequency and 3db cutoff increased with increasing distances from the pial surface (Fig. 3b-e). Input resistance was lowest for neurons found in deep L3, whereas resting membrane potential and sag increased with distance from the pial surface, with deeper neurons appearing more depolarized (Fig. 3f). Except for small differences in resting potential and 3dB cutoff, there were no significant differences in HCN dependent properties between acute and culture slice preparations, nor was the slope of the depth dependence different between preparations (Supplementary Table 3). Furthermore, these trends were apparent and conserved in recordings obtained from individual animals (Fig. S4). Together these data demonstrate that HCN-dependent properties in temporal cortex are enriched in supragranular pyramidal neurons found farther from the pial surface in macaques, even within individual animals, like in human MTG^14,22,23^.

Next, we tested whether the depth-dependence was present in neocortical areas outside of temporal cortex using a similar approach in primary motor cortex, a region rarely available in human surgical resections. In general, HCN-related properties in motor cortex neurons were less dependent on the distance from the pial surface compared with temporal cortex. In motor cortex 3dB cutoff was lower in deeper compared with superficial neurons and there was no difference in resonance frequency across the cortical depth (Fig. S5; Supplementary Table 4). Although sag increased with depth from the pial surface, input resistance and resting potential were constant (Fig. S5; Supplementary Table 4). Thus, although prominent HCN related membrane properties are observed in supragranular pyramidal neurons in both temporal and motor cortical areas, depth from pia dependent increases may be specific to certain neocortical areas and/or cell types.

### Blocking HCN channels decreases resting conductance and eliminates band pass filtering properties in macaque supragranular neurons

To test whether HCN channels contribute to the resting conductance and subthreshold filtering properties of macaque supragranular pyramidal neurons, we bath applied the HCN channel blocker ZD7288 (10 μM) and measured changes in subthreshold membrane properties. For these experiments we targeted neurons throughout the depth of layer 2/3 to assess whether blocking HCN conductance has a larger effect on neurons located deeper with layer 3, where HCN properties were most pronounced. For each recorded neuron, sag ratio, input resistance and resonance frequency were measured before and after the application of ZD7288, at a common membrane potential of-65 mV.

Blocking HCN channels hyperpolarized the resting membrane potential by 5.34 +/-1.44 mV (Fig. S6, pre =-66.17 +-0.48, post =-71.51 +-1.45; p=1.6e-03, paired t-test), eliminated sag (Fig. 4b, Fig. S6, pre = 1.24 +-0.04, post = 1.04 +-0.01; p=5.8e-05, paired t-test) and eliminated membrane resonance (pre =2.03 +-0.27, post = 0.48+-0.03; p=4.2e-05, paired t-test) in response to the chirp stimulus (Fig 4b,e). Additionally, ZD7288 increased input resistance by 56.69+/-8.94% (Fig. 4c, pre = 86.88+-11.09, post = 126.34 +-10.46; p=3.0e-07, paired t-test), indicating that HCN channels contribute substantially to the resting conductance of macaque supragranular pyramidal neurons in TCx. The effect of blocking HCN channels on input resistance was larger for neurons found farther from the pial surface (Spearmann’s rho=0.47, p=0.043; Fig. 4d), but the change in resonance (Spearmann’s rho = 0.07, p = 0.77), resting potential (Spearmann’s rho = 0.11, p = 0.66) and sag (Spearmann’s rho = 0.24, p = 0.31) did not depend on depth from pia (Fig. S6). Notably, the change in resonance (Spearmann’s rho = 0.64, p = 0.003) and sag (Spearmann’s rho = 0.71, p = 0.0006) correlated strongly with the change in input resistance (Fig. 4f; Fig. S6). Taken together, these findings provide pharmacological evidence that HCN channels contribute to the physiological properties of macaque supragranular pyramidal neurons and suggest that the neurons with the most pronounced HCN-dependent properties also have the most HCN-conductance, regardless of their position within L2/3.

**Figure 4.**
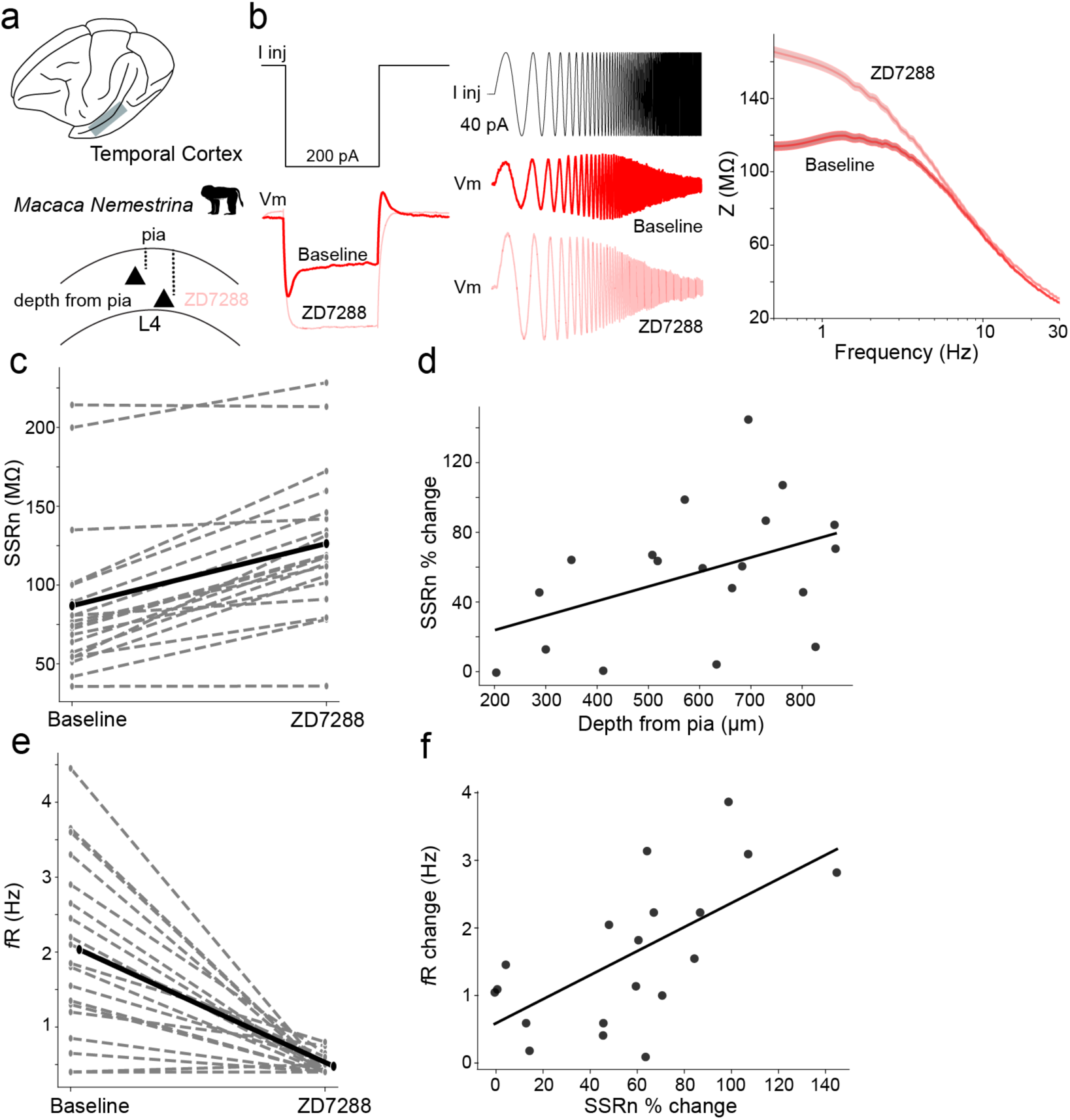
Blocking HCN conductance eliminates subthreshold resonance and decreases membrane conductance in supragranular neurons in temporal cortex. a) We measured the effect of ZD7288 on HCN channel-related properties of supragranular neurons in the temporal cortex of *M.nemestrina.* b) *Left* - Hyperpolarizing current injection step and example membrane response of a cell before and after ZD7288 application. *middle* - Chirp current injection and example membrane response of a cell before and after the bath application of ZD7288. *Right* - average impedance amplitude profile of all cells (n = 19) before and after ZD7288 application. c) Input resistance for each neuron (dashed lines) before and after ZD7288 application. Mean values are connected by the thick line. d) Percent change in input resistance plotted as a function of distance from the pial surface. e) Resonance frequency for each cell before and after ZD7288 application. f) Change in resonance frequency plotted as a function of percent change in input resistance.

### Conserved and divergent HCN-dependent membrane properties in human versus macaque cell types

Although these data suggest that HCN channels contribute significantly to the subthreshold membrane properties of supragranular pyramidal neurons in non-human primates, cross-species differences may exist as has been reported for L5 ET neuron physiology ^19^. To directly compare HCN-dependent membrane properties in human versus macaque neocortex, we utilized a previously published human supragranular glutamatergic Patch-seq dataset^10^, which we supplemented with additional new samples. To compare homologous cell types, we mapped the human and macaque Patch-seq datasets to a consensus MTG taxonomy derived from snRNA-seq across several non-human primate species (Figure 5a,b) ^41^, which contains three types of supragranular glutamatergic neurons, L2/3 IT_1-3. Samples mapping to L2/3 IT_3 were enriched in the upper portion of layer 2/3 whereas samples mapping to L2/3 IT_1 were found throughout the depth of layer 2/3 and we rarely sampled from L2/3 IT_2 (Figure 5b). This spatial distribution matched the distribution of these cell types in a multiplexed error-robust fluorescence in situ hybridization (MERFISH) dataset from human MTG mapped to the same reference taxonomy (Fig. S2). Notably, *HCN1* expression increased as function of somatic depth from pia in this dataset (Fig. S2). Because we rarely sampled neurons mapping to L2/3 IT_2, we restricted subsequent analysis to L2/3 IT_1 and L2/3 IT_3.

**Figure 5.**
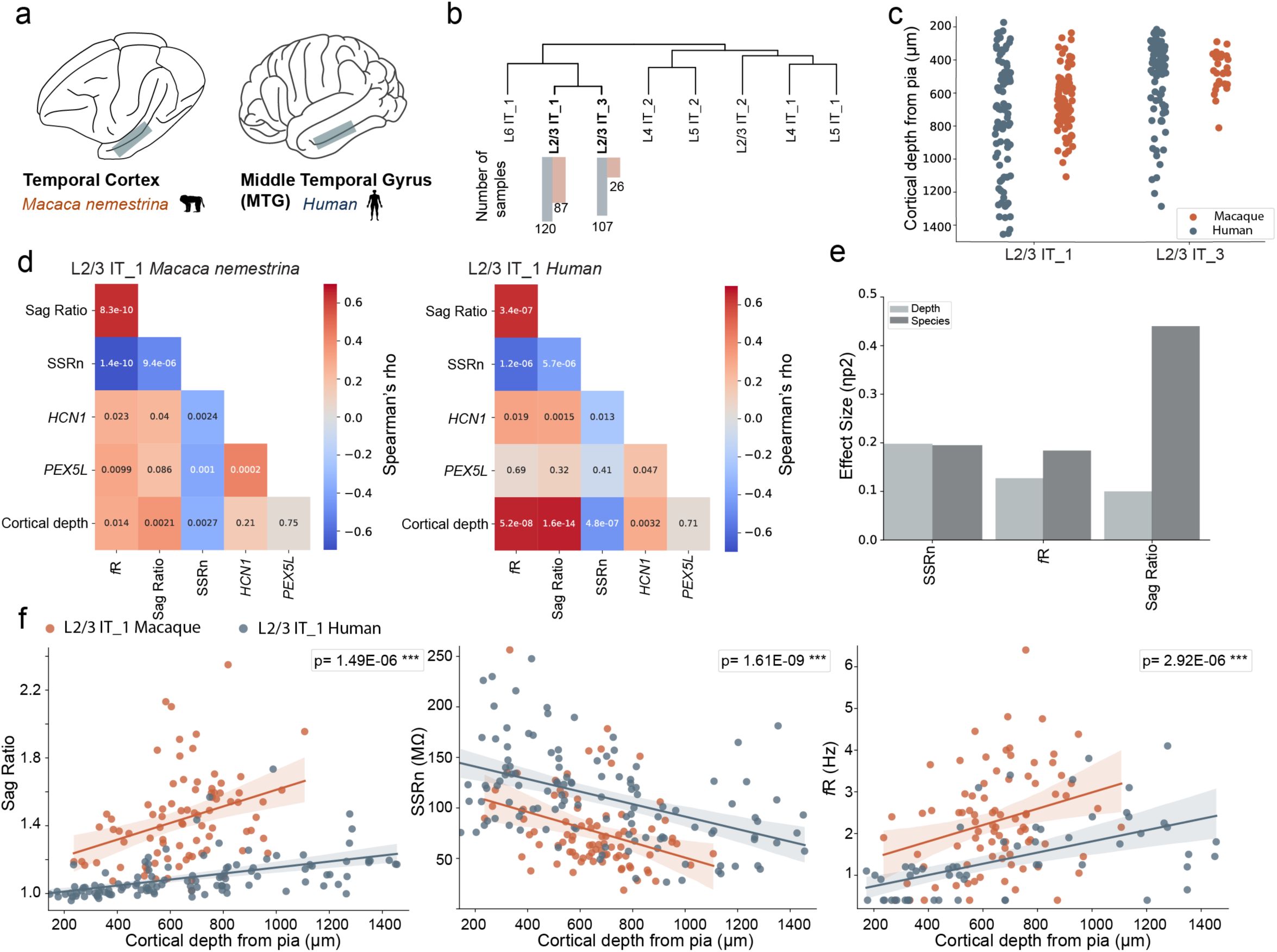
Patch-seq reveals divergence and similarities in HCN-dependent properties in temporal cortex of macaque versus human. a) Patch-seq samples were collected from macaque (*M. nemestrina*) and human temporal cortex. b) Dendrogram showing that samples predominately mapped to two clusters in the cross-species consensus taxonomy. The bar graphs denote the number of samples mapping to each cluster. c) Depth from pia of each Patch-seq sample by transcriptomic type. d) Spearman’s rho correlations between HCN channel-dependent membrane properties, depth from pia and HCN channel-gene expression for human and macaque samples mapping to L2/3 IT_1. e) Depth from pia and species effect sizes for HCN channel dependent features. f) HCN channel-dependent properties plotted as a function of distance from pia for each species. p values denote significance for depth from pia effect (Supplementary Table 5).

We first asked whether the HCN-dependent membrane properties correlated with depth from pia for both L2/3 IT_1 and L2/3 IT_3 neurons of human and macaque. Additionally, gene expression obtained during Patch-seq enabled us to address whether HCN-related gene expression correlated with depth from pia and HCN-dependent membrane properties. To this end, we performed pairwise correlations of HCN-dependent properties (*f*_R_, sag, input resistance) HCN-related gene expression (*HCN1 and PEX5L*) and depth from pia. HCN-related membrane properties correlated with depth from pia in L2/3 IT_1 in both species (Figure 5d), whereas in L2/3 IT_3, correlation coefficients were either not significant or were smaller (Fig. S7).

Additionally, *HCN1* gene expression correlated with sag, input resistance and resonance in L2/3 IT_1 in both species and individual HCN-related membrane properties correlated strongly with each other (Fig. 5d). Thus, individual HCN-related properties correlate with each other, with *HCN1* gene expression and with depth from pia within a single transcriptomic cell type in both human and macaque temporal cortex.

To directly test for cross-species differences within these cell types, we fit each HCN-dependent property with multiple linear regression in which depth from pia and species were the independent variables. Although HCN-dependent properties varied with depth from pia for neurons mapping to L2/3 IT_1 in both species (Figure 5e, f; Supplementary Table 5; 2-Way ANOVA, effect of depth from pia on sag ratio, p= 1.49e-06; on input resistance, p= 1.61e-09; on resonance frequency, p = 2.92e-06), there were notable cross-species differences as well.

Sag and resonance frequency were higher, and input resistance was lower in macaque compared with human L2/3 IT_1 neurons (Fig.5e,f, Supplementary Table 5; 2-Way ANOVA, effect of species on sag ratio, p= 1.87e-28; on input resistance, p= 4.13e-09; on resonance frequency, p= 1.49e-07). Additionally, the slope of the relationship between sag and depth from pia was steeper in macaque compared with human L2/3 IT_1 neurons (Fig.5e,f, species by depth from pia interaction, p= 3.40e-02). These data demonstrate that within the L2/3 IT_1 type, HCN-dependent properties vary with depth from pia in both human and macaque temporal cortex. Additionally, HCN-dependent properties for this specific cell type are more pronounced in macaque.

In contrast, we did not observe differences between macaque and human for the L2/3 IT_3 type for most intrinsic features analyzed. Here, the HCN-dependent properties were mildly dependent on cortical depth (Fig.S7; Supplementary Table 5; 2-Way ANOVA, Effect of depth from pia on sag ratio, p= 1.2e-02; on input resistance, p= 7.5e-03; on resonance frequency, p= 7.5e-03). Except for sag ratio, HCN-related properties were not significantly different between humans and macaques for the L2/3 IT_3 cell type. Moreover, the effect size of both species and cortical depth was low for all intrinsic features considered (Fig.S7; Supplementary Table 5; 2-Way ANOVA, Effect of species on sag ratio, p= 4.37e-03; on input resistance, p= 0.96; on resonance frequency, p= 0.66). These data highlighted that cross-species differences in the biophysical properties of supragranular neurons depend upon the neuron subtype.

## DISCUSSION

Human supragranular pyramidal neurons display enriched HCN channel-related gene expression and intrinsic membrane properties compared with their counterparts in rodent neocortex ^10,22^. Here, we provided evidence that these differences in supragranular neuron physiology reflect broader distinctions between primate and rodent neurons. *HCN1* subunit expression in primates was roughly equivalent in supragranular glutamatergic and L5 ET neurons, a cell population known for enriched HCN-dependent properties ^52^. HCN-dependent intrinsic membrane properties were ubiquitous in supragranular pyramidal neurons in motor and temporal cortex of both the New World and Old World monkey species we studied. In the temporal cortex of pig-tailed macaques, we found that HCN-dependent properties were most pronounced in neurons located farther from the pial surface, like what has been reported in human MTG ^10,22^. Using Patch-seq, we showed that this depth from pia-dependent variation in HCN properties was most robust in the L2/3 IT_1 cell type in both human and macaque neocortex. For this cell type, HCN-dependent properties were generally more pronounced in macaque compared with human neurons. Finally, we did not observe strong cross-species differences in HCN properties in the other major supragranular glutamatergic cell type, L2/3 IT_3, suggesting that species differences are cell type-dependent.

### Functional implications of HCN channel expression in primate supragranular pyramidal neurons

In primates, the supragranular layers comprise approximately 46% of the total thickness of the neocortex compared with only 19% in rodents ^1^. Consequently, the total dendritic length of primate supragranular pyramidal neurons is longer than in rodent neurons ^10,12,53^ (but see ^11^). In a passive neuron, the capacitive filtering imposed by the cable properties of such long dendrites would be severe and the temporal window for synaptic integration would heavily depend upon the site of synaptic input ^54^. HCN channel expression counteracts this capacitive filtering in several neuron types, including in human supragranular pyramidal neurons, thus ensuring that the kinetics of synaptic potentials at the soma are relatively independent of the site of origin at the dendrite ^12,22,29,30,46^. This helps ensure that the temporal summation of synaptic input at the site of action potential generation does not depend on the site of synaptic input. We suggest that HCN channel expression serves a similar function in non-human primate supragranular pyramidal neurons.

In addition to counteracting capacitive filtering, HCN channel expression can promote the subthreshold integration of inputs containing certain frequencies. In a previous study, we provided evidence that in human supragranular pyramidal neurons, HCN conductance promotes the integration of inputs containing delta/theta band frequencies ^22^. Thus, subthreshold integration in non-human primate supragranular pyramidal neurons may also be tuned to inputs in the delta/theta frequency range. In this regard, the conserved expression of HCN channels in supragranular pyramidal neurons amongst primates may contribute to the conserved spectral signatures observed in local field potential recordings in human and non-human primate neocortical areas. In the supragranular layers of primate species there is a prominent delta/theta (1-8 Hz) band in the local field potential that is absent in the supragranular layers of mouse neocortex ^55^. We propose that the ubiquitous expression of HCN channels in supragranular neurons in primates contributes to these spectral signatures in primate neocortex *in vivo*.

Conversely, the absence of HCN channel expression in rodents may contribute to the lack of prominent delta/theta frequency bands in the supragranular layers *in vivo*.

Additionally, the prominent HCN conductance observed in primates could provide a mechanism for fine-tuned regulation of synaptic integration in species with more complex cognitive functions, compared to rodents. In this context, HCN channels represent key targets of both neuromodulation and intrinsic plasticity, which regulate channel voltage dependence, gating kinetics, and consequently, neuronal and network excitability ^46,56,57^.

### Cell type and species differences in HCN channel-related properties

We found that HCN channel-dependent properties were ubiquitous amongst primate species, but we observed notable cross-species and cross-areal differences. For example, resonance frequency was higher in motor cortex than temporal cortex. This is consistent with previous observations that HCN channel-dependent properties are more pronounced in L5 ET neurons of motor cortex versus temporal cortex of the macaque ^45^. Thus, there may be general cross-areal differences in HCN-dependent properties in the primate neocortex in supragranular layers as well. Alternatively, differences in the cell type composition between neocortical areas ^8,58^ may contribute to the cross-areal differences we observed in HCN channel properties. For instance, there may be cell types enriched in motor, but not temporal areas with especially pronounced HCN-dependent properties. This hypothesis could soon be directly tested by mapping our Patch-seq dataset to forthcoming macaque snRNA-sequencing based taxonomies of M1 produced through the BRAIN Initiative.

In addition to cross-areal differences within the macaque neocortex, we also observed cross-species differences between human and macaque neurons, especially within the L23_IT1 cell type. Notably, L23_IT1 macaque neurons had more pronounced HCN-dependent properties and had a lower input resistance. Thus, despite presumably being larger than macaque neurons, human neurons possess a higher input resistance, suggesting that resting conductance is lower in human neurons than in macaque. This is consistent with a previous study demonstrating that L5 human neurons exhibit ionic conductance densities that are lower than expected based on their size ^19^. Thus, if L23_IT1 neurons display a similar allometric scaling rule that has been described for L5 neurons, human neurons would be outliers.

Notably, the cross-species differences we observed in the L2/3 IT_1 cell type were largely absent in the L2/3 IT_3 type, suggesting that innovations in human neuron properties are cell type specific.

Although addressing the mechanisms underlying species differences will require additional investigation, we offer a few speculative possibilities. Species differences in the HCN channel-dependent properties may reflect differences in dendritic morphology between species. In this context, the cross-species similarity/dissimilarity of the HCN channel properties we found could reflect similarity/dissimilarity in dendritic morphology within specific cell types. Additionally, cell type and/or species-specific differences in HCN channel-related protein isoforms may contribute to our findings, as it has been described for Na+ channel function^59^. For example, there are several isoforms of TRIP8B, each of which has a different effect on the surface expression or voltage-dependent properties of HCN channels and associated subthreshold membrane properties. Although we observed ubiquitous expression of the gene encoding TRIP8B (*PEX5L*) in human and macaque neurons, cross species differences in alternative splicing and isoform expression within the L2/3 IT_1 type may contribute to our observations.

### Study limitations

Our recordings were performed in current clamp and were restricted to the soma. Therefore, our conclusions are applicable to the perisomatic compartment only and do not address potential species differences in channel properties. Future studies utilizing direct measurements of HCN conductance in voltage clamp recordings may reveal the subcellular localization of HCN channels and differences in channel kinetics. Finally, our measurements were restricted to subthreshold membrane properties. Cell type and species differences in suprathreshold membrane properties associated with HCN channels and other ionic conductance could contribute to substantial differences in neuron function. Nonetheless we provide evidence of strong HCN-dependent properties in supragranular pyramidal neurons in non-human primates, supporting the hypothesis that the presence of HCN conductance is conserved across primate species and does not represent a human specialization. Our approach of using Patch-seq across various mammalian species serves as a roadmap for understanding human and primate specializations.

## Acknowledgments

We thank the Allen Institute for Brain Science Tissue Procurement, Processing and Facilities teams for their help in coordinating logistics of human surgical tissue collection, transport and processing. We thank Tom Reh for generously donating squirrel monkey brain tissue. We thank the Washington National Primate Research Center veterinary support staff and Chris English for support during tissue procurement. The WaNPRC is supported by the NIH Office of Research Infrastructure Programs (ORIP) under award number P51OD010425 and U420D011123. We thank the Viral technology team for packaging AAV-based tools. We are grateful to Dana Rocha, Anish Chakka, Michael Tieu, Christine Rimorin, Delissa McMillen, and Trangthanh Cardenas for processing the Patch-seq samples. We thank Christina Alice Pom, Stuard Barta, Alana Oyama, Angela Ayala, and Jeanelle Ariza for biocytin staining. We are grateful to Naomi Martin, Nasmil Valera Cuevas, Paul Olsen, Jazmin Campos, Josh Nagra, Emily Gelfand, and Delissa McMillen for the generation of the MERFISH data. We thank Tim Jarsky for assistance related to the MIES acquisition software. We are grateful to Jack Waters, Lydia Potekhina, Shea Ransford, Zoe Juneau and Sam Hastings for imaging biocytin filled neurons. A special thanks to Andres Barria (University of Washington Department of Neurobiology & Biophysics) for helping with brain slice culturing techniques. We thank Natalia Goriounova and Huibert Mansvelder for helpful discussions on figures and analysis. Finally, we wish to thank Paul G. Allen for his vision, encouragement, and support.

This research has been supported by two primary U.S. National Institutes of Health (NIH) grants: R01NS123959 and UG3MH120095. The contents of the present study are solely the responsibility of the authors and do not necessarily represent the official view of the NIH, WaNPRC, ORIP or NCATS.

## Conflict of interest

The authors declare no competing financial interests

## Author contributions

Conceptualization, B.E.K., N.C.D, J.T.T.; software, N.C.D., S.S., and B.E.K.; data curation, B.E.K., C.R., N.C.D., A.L.K., B.L.G., J.S.H., A.P.P., C.C.; resources, B.E.K., N.C.D., J.T.T., C.R., S.S., X.O.-A., M.H., N.T., M.S., A.L.K., B.L.G., J.S.H., A.P.P., C.C.; formal analysis, B.E.K., C.R., X-P.L, N.C.D., and M.S.; supervision, B.E.K., N.C.D., J.T.T. and W.J.S; funding acquisition, B.E.K., N.C.D., J.T.T., W.J.S. and E.L.; validation, N.C.D., S.S., B.E.K.; investigation, C.R., B.E.K., N.C.D. and S.S.; visualization, C.R. and B.E.K.; methodology, B.E.K., N.C.D., J.T.T., X.O.-A., M.H., J.T.T., T.B.; writing - original draft, C.R., B.E.K.; writing – review and editing, B.E.K., C.R., N.C.D., X-P.L. and J.T.T.; project administration, J.T.T., N.C.D. and B.E.K.

## Notes

### Competing Interest Statement

The authors have declared no competing interest.

