## Supplemental figures and tables for "HCN channels reveal conserved and divergent physiology in supragranular pyramidal neurons in primate species"

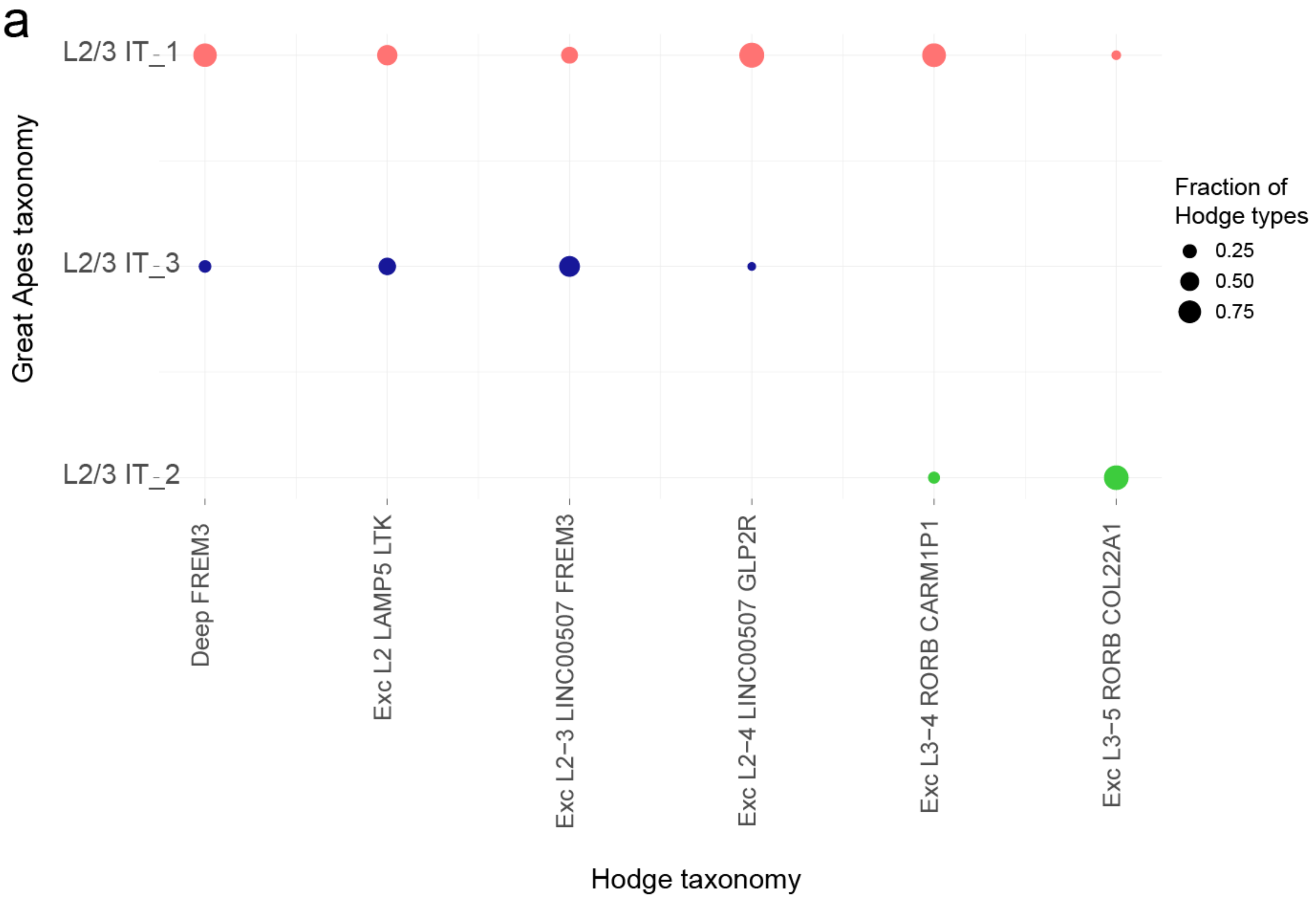


**Supplementary Figure 1 Matching the Hodge et al., MTG taxonomy with the Great Apes consensus taxonomy** a) Dotplot of human Patch-seq samples mapped against the Hodge[6] taxonomy versus the Great Apes consensus Taxonomy[41].

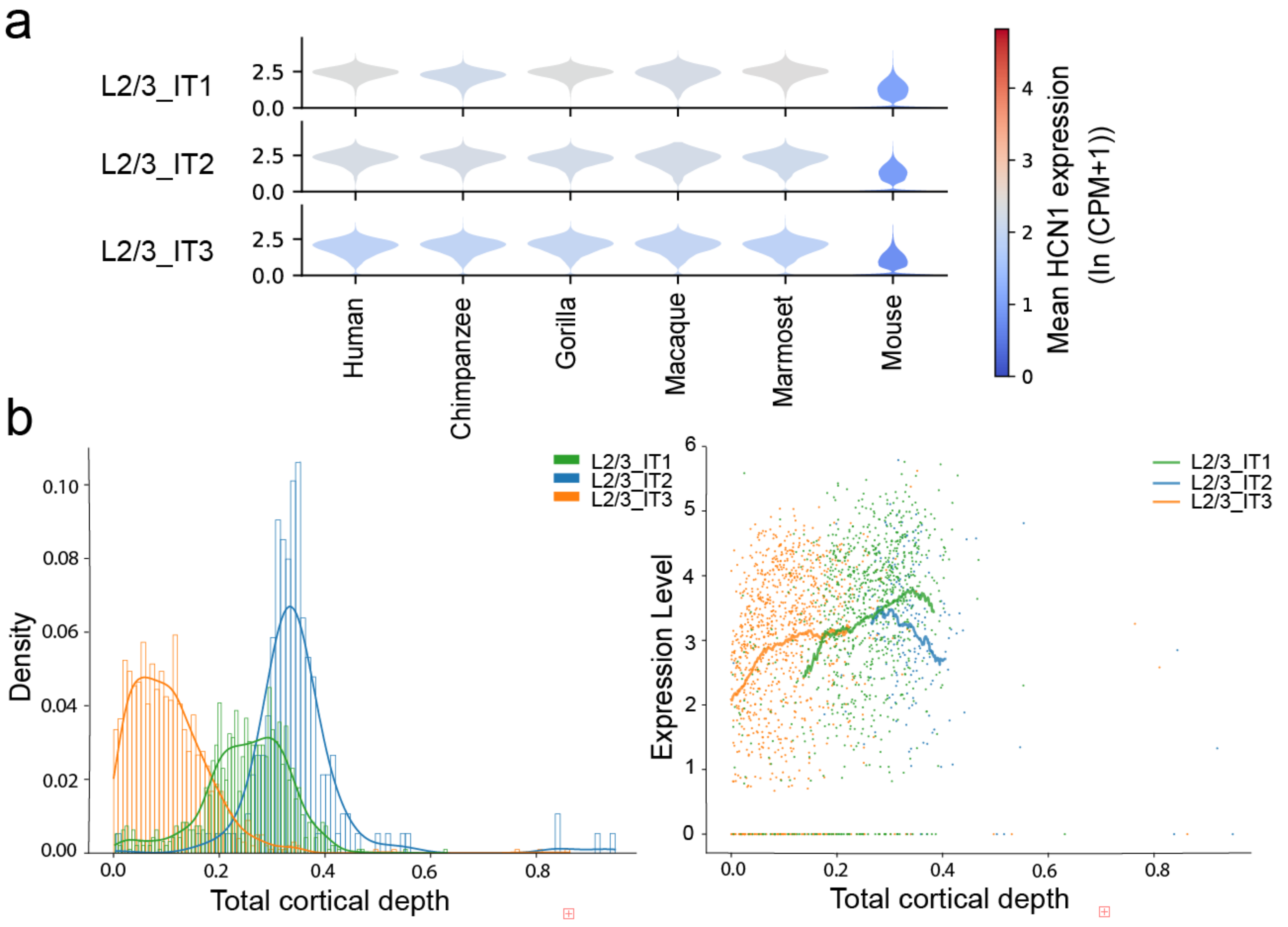


**Supplementary Figure 2 – HCN channel related gene expression analysis across species.** a) Violin plot of *HCN1* expression in each type of L2/3 IT neuron in the consensus great apes MTG taxonomy across six species: mouse, marmoset, macaque, gorilla, chimpanzee and human. b) Spatial distribution of the L2/3 IT types in human MTG revealed by MERFISH. c) *HCN1* expression as a function of cortical depth for L2/3 IT t-types in human MTG, as revealed by MERFISH.

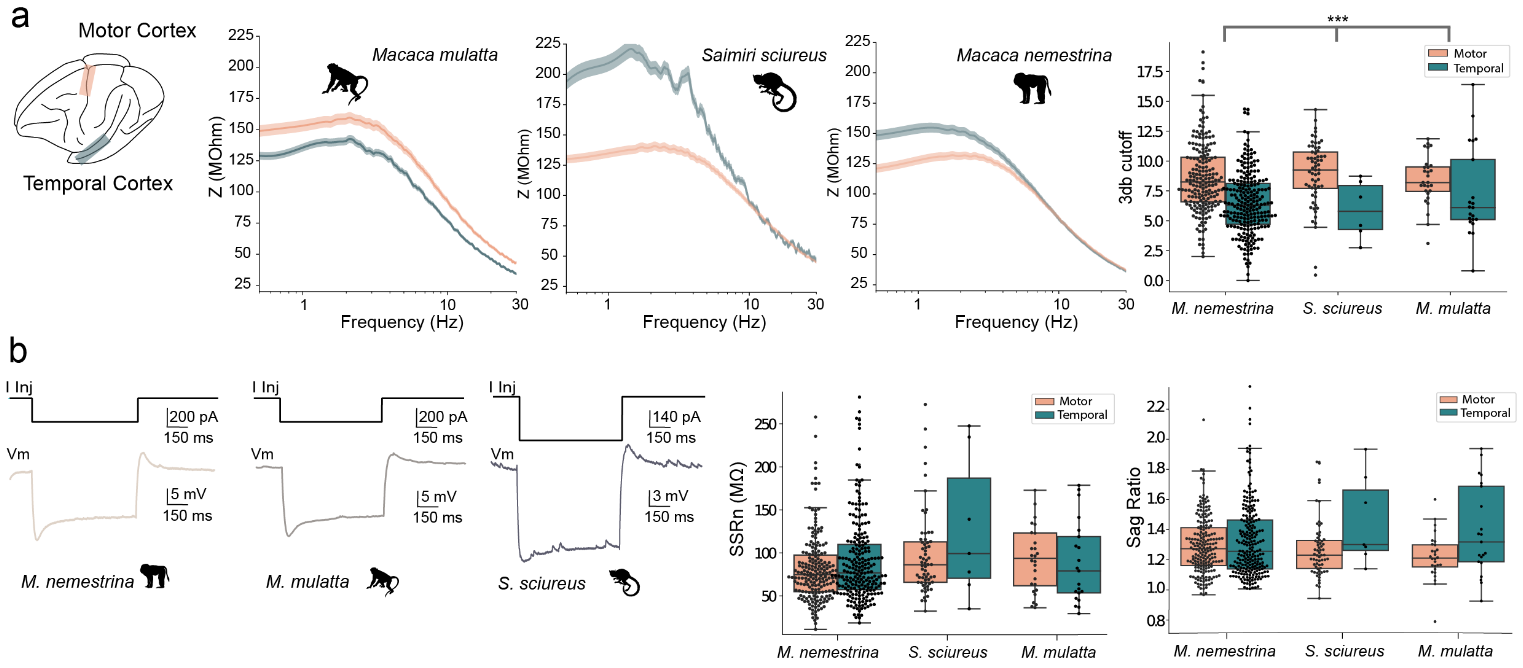


**Supplementary Figure 3 – Additional HCN dependent properties in non-human primate species and cortical areas.** a) Average (solid line) impedance amplitude profiles for neurons recorded in the temporal cortex and motor cortex of *Macaca nemestrina*, *Macaca mulatta* and *Saimiri sciureus,* and 3dB cutoff frequency for each species and neocortical area. Shaded area corresponds to SEM. b) Examples responses to hyperpolarizing current injection in each species. Input resistance and sag ratio for each species and neocortical area. *** denotes significant effect of area. See Supplementary Table 2 for statical comparisons between groups.

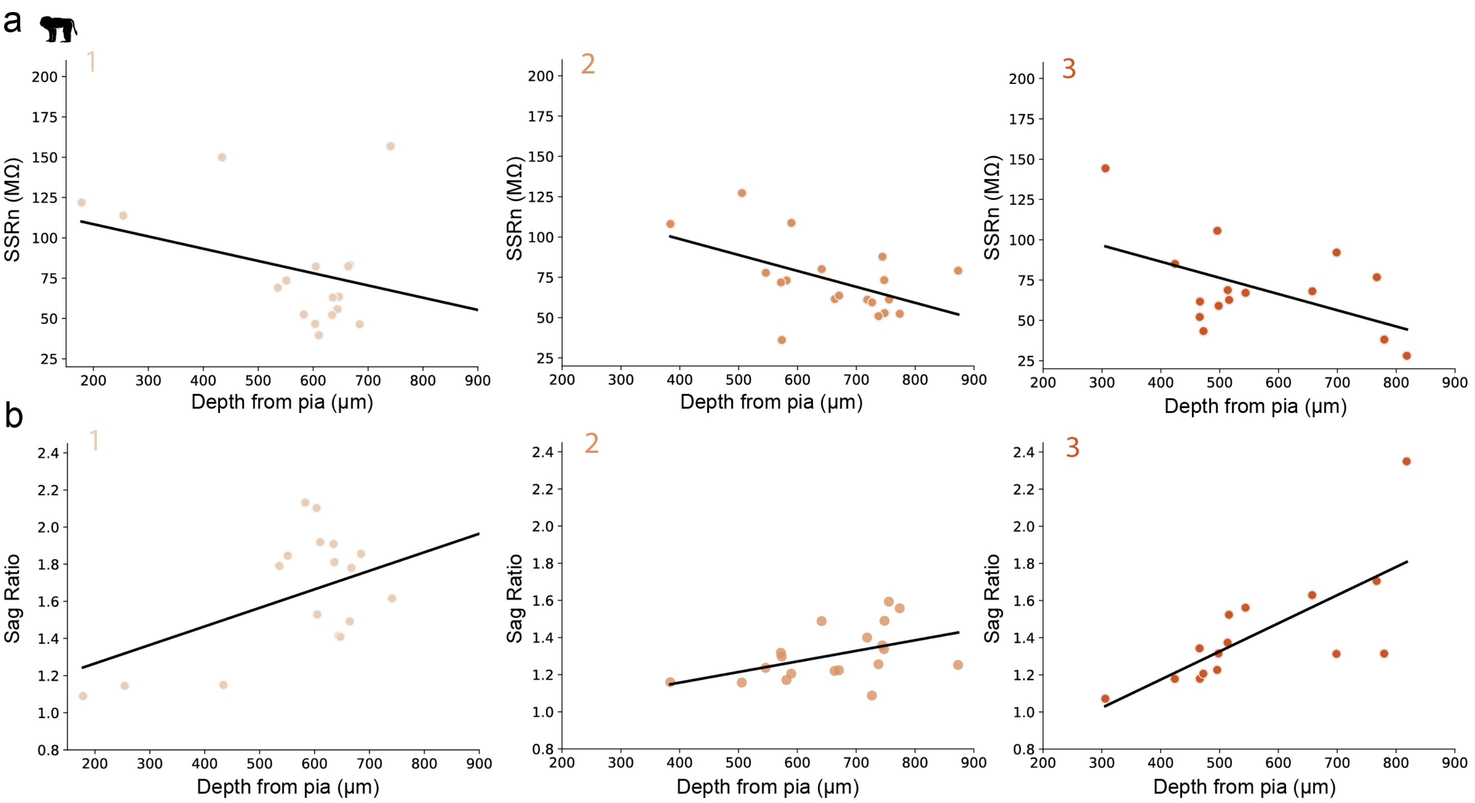


**Supplementary Figure 4 – Depth from pia dependent properties in single subjects.** a) Input resistance and b) sag ratio as a function of depth from pia in individual pig-tailed macaques.

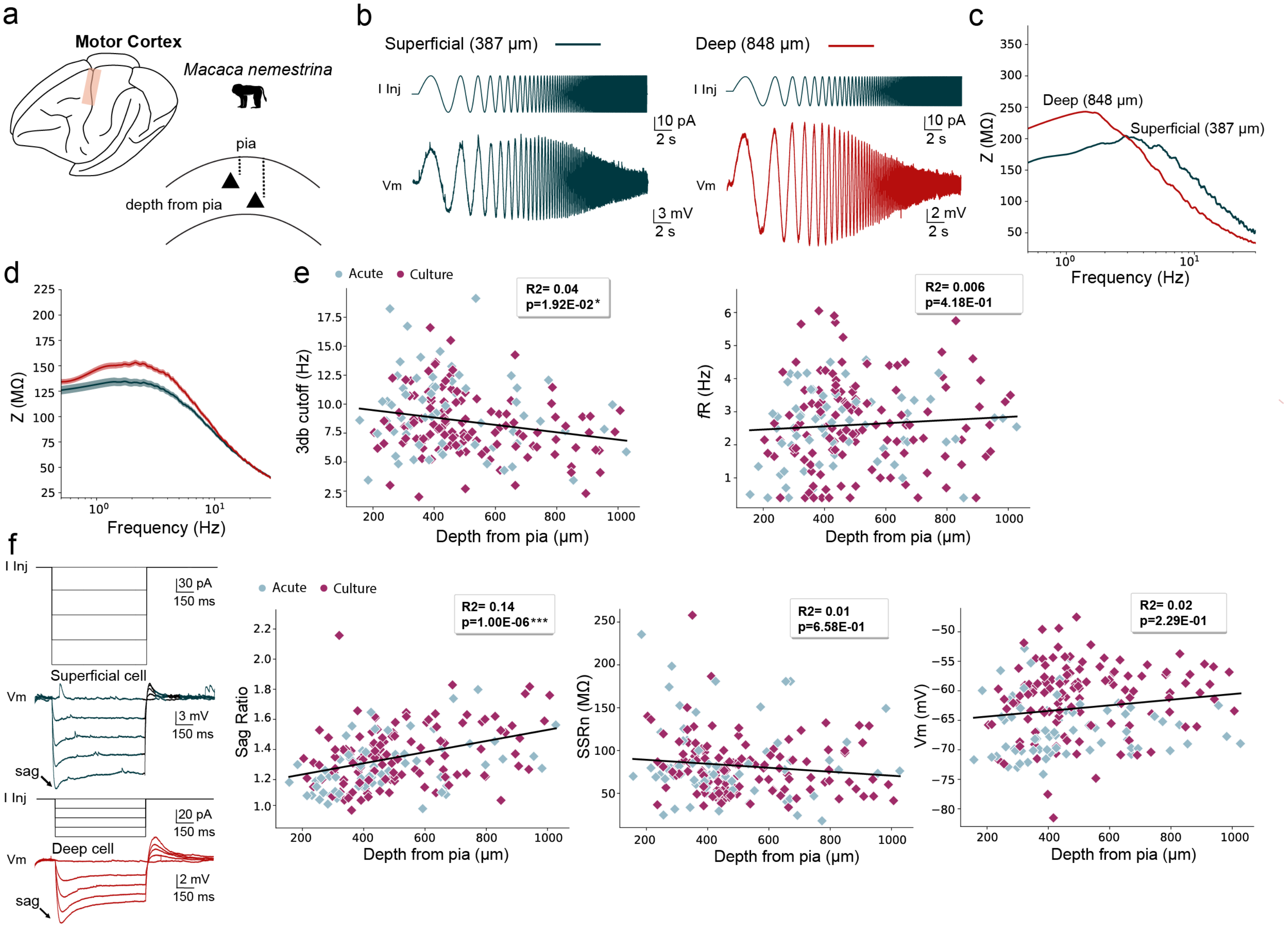


**Supplementary Figure 5 – HCN channel-related intrinsic membrane properties across the radial axis of the motor cortex of the pig-tailed macaque.** a) We quantified the subthreshold features of supragranular pyramidal neurons throughout the radial depth of the motor cortex in *Macaca nemestrina.* b) example voltage response to a chirp stimulus for a superficial and deep neuron and c) corresponding impedance amplitude profiles. d) impedance amplitude profiles averaged across the superficial most one third (red) and deepest one third of neurons (green). e) 3dB cutoff and resonance frequency plotted as a function of distance from pial surface. f) Example membrane response to a series of hyperpolarizing current injections for a superficial and deep neuron. g) Sag ratio, steady-state input resistance and resting potential plotted as a function of distance from the pial surface (2-way ANOVA, Supplementary Table 4).

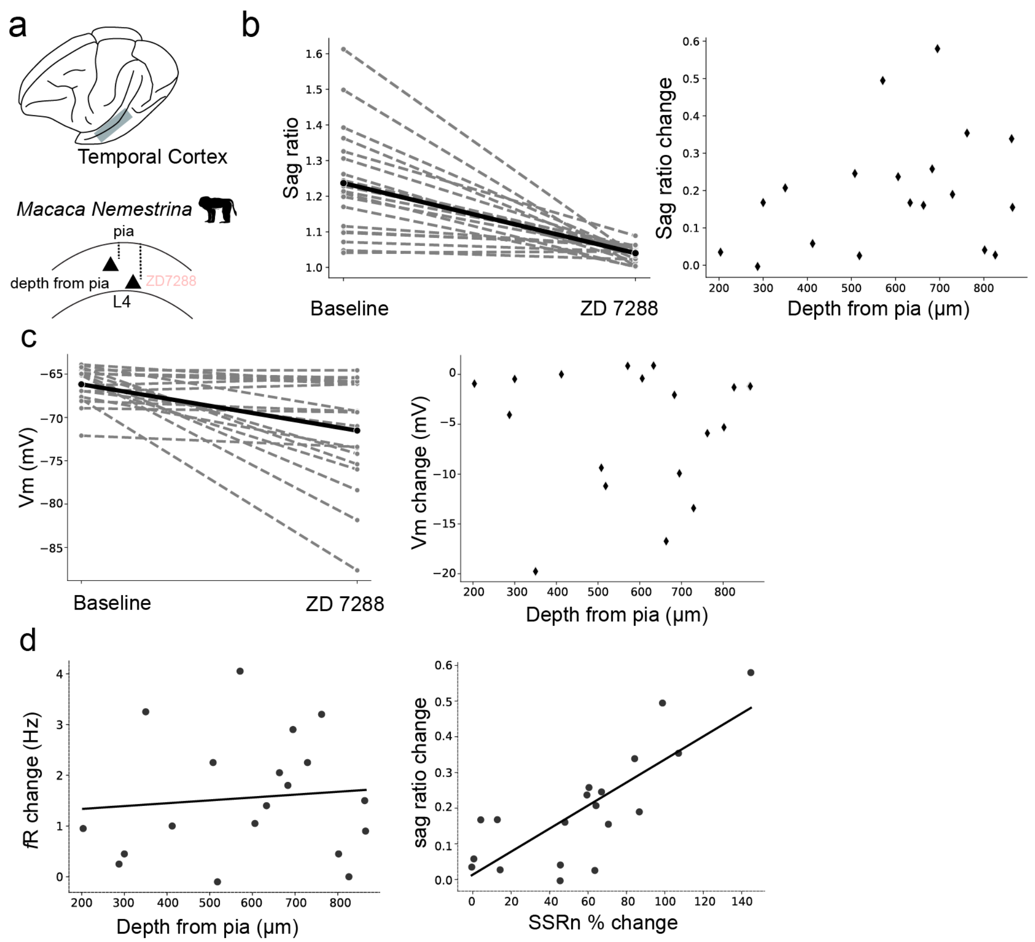


**Supplementary Figure 6 - Blocking HCN conductance decreases sag ratio and depolarizes the resting membrane potential in supragranular neurons in temporal cortex.** a) Dataset restricted to supragranular layers of *M.nemestrina* in temporal cortex. b) *Left* – Sag ratio for each neuron (dashed lines) before and after ZD7288 application. Mean values are connected by the thick line. *Right* - Change in *s*ag ratio plotted as a function of distance from the pial surface. c) Left– Resting membrane potential for each cell before and after ZD7288 application. Right – Change in resting membrane potential plotted as a function of depth from the pial surface. d) Absolute change in resonance frequency as a function of depth from pia. *e)* Absolute change in sag ratio as a function of change in input resistance.

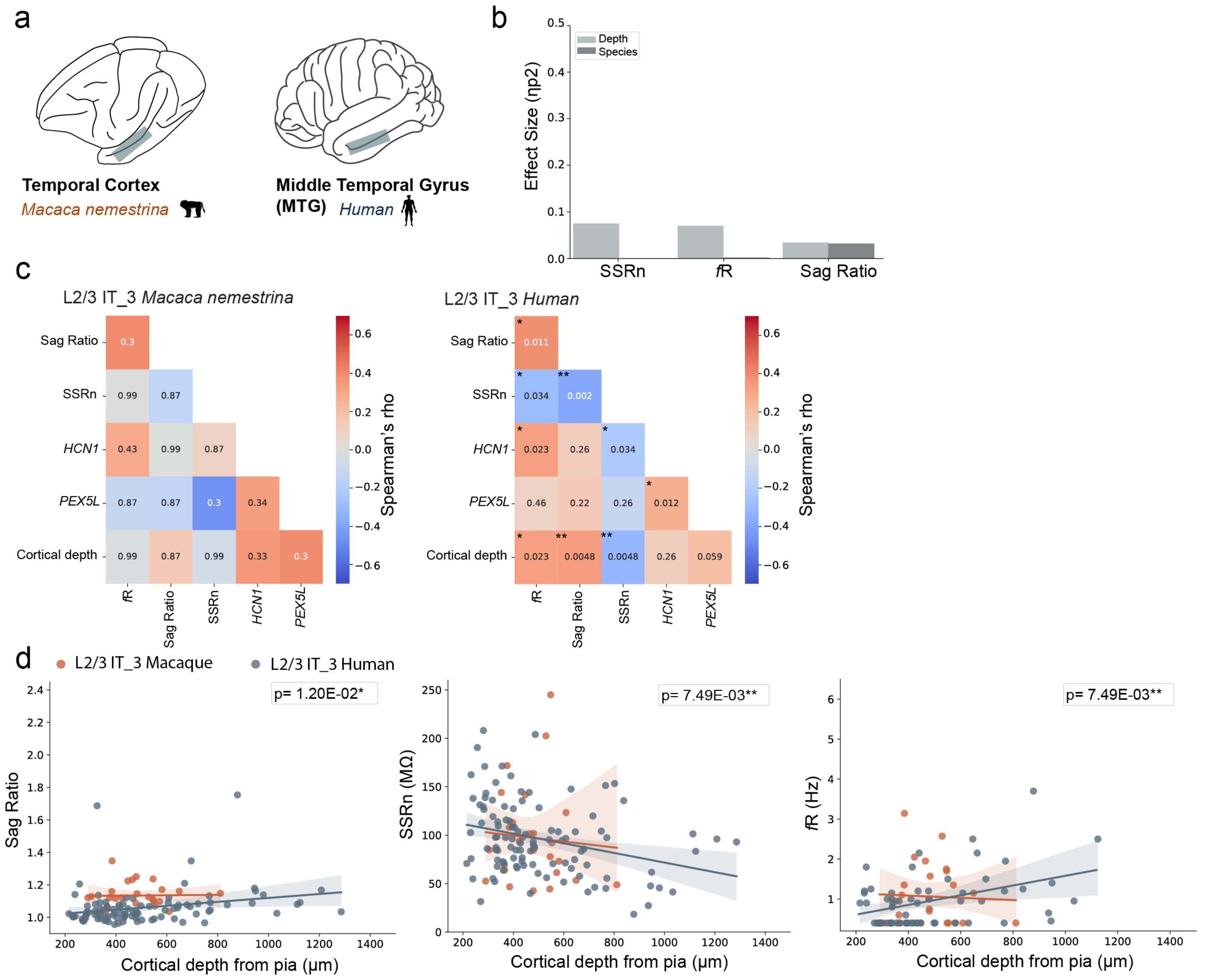


**Supplementary Figure 7 - Cross-species comparison of the L2/3 IT_3 cluster.** a) Dataset restricted to supragranular temporal cortex neurons in *Macaca nemestrina* and supragranular middle temporal gyrus neurons in *Human.* b) Depth from pia and species effect sizes for HCN channel dependent features in L2/3 IT_3 neurons. c) Spearman's rho correlations between HCN channel-dependent membrane properties, depth from pia and HCN channel-gene expression for human and macaque samples mapping to L2/3 IT_3. d) HCN channel-dependent properties plotted as a function of distance from pia for each species for the L2/3 IT_3 cluster. P values are for depth from pia effect (Supplementary Table 5).

**Demographic data for Human Donors**

| Age (years) | Sex | Seizure history (yrs) | Cause for resection | Resection location |
|---|---|---|---|---|
| 42 | M | 14 | Epilepsy | Left Temporal |
| 37 | F | NA | Epilepsy | Left Temporal |
| 64 | M | 1 | Tumor | Right Temporal |
| 18 | F | 2 | Epilepsy | Left Temporal |
| 75 | F | NA | Epilepsy | Right Temporal |
| 58 | F | NA | Epilepsy | Right Temporal |
| 24 | F | 6 | Epilepsy | Left Temporal |
| 68 | M | NA | Tumor | Right Temporal |
| 72 | F | NA | Tumor | Right Temporal |
| 41 | M | 5 | Epilepsy | Left Temporal |
| 29 | M | 4 | Epilepsy | Right Temporal |
| 19 | F | 18 | Epilepsy | Left Temporal |
| 45 | M | NA | Epilepsy | Right Posterior Temporal |
| 24 | M | NA | Tumor | Left Temporal |
| 32 | F | 3 | Tumor | Right Temporal |
| 24 | M | 5 | Epilepsy | Right Anterior Temporal |
| 32 | M | NA | Tumor | Right Temporal |
| 28 | M | 18 | Epilepsy | Right Temporal |
| 71 | M | NA | Tumor | Left Temporal |
| 53 | M | NA | Tumor | Left Temporal |
| 54 | M | NA | Tumor | Right Temporal |
| 20 | F | 2 | Epilepsy | Right Temporal |

Supplemental Table 1

| Variable | Bejamini-Hochberg adjusted p-value | variance explained (η2) |
|---|---|---|
| *Species* | | |
| sag ratio | 8.82E-01 | 0.00 |
| input resistance (SSRn) | 1.03E-01 | 0.01 |
| resonance frequency (fR) | 6.26E-01 | 0.00 |
| 3db cutoff | 8.82E-01 | 0.00 |
| *Preparation* | | |
| sag ratio | **5.88E-04** | 0.03 |
| input resistance (SSRn) | 6.93E-01 | 0.00 |
| resonance frequency (fR) | 2.48E-01 | 0.00 |
| 3db cutoff | **9.14E-03** | 0.01 |
| *Cortical area* | | |
| sag ratio | 1.07E-01 | 0.01 |
| input resistance (SSRn) | 1.33E-01 | 0.00 |
| resonance frequency (fR) | **5.82E-07** | 0.05 |
| 3db cutoff | **1.01E-14** | 0.11 |
| *Cortical area:Preparation* | | |
| sag ratio | 8.42E-01 | 0.00 |
| input resistance (SSRn) | 8.42E-01 | 0.00 |
| resonance frequency (fR) | 8.42E-01 | 0.00 |
| 3db cutoff | 8.42E-01 | 0.00 |
| *Preparation:Species* | | |
| sag ratio | 3.14E-01 | 0.00 |
| input resistance (SSRn) | 2.96E-01 | 0.01 |
| resonance frequency (fR) | 2.28E-01 | 0.01 |
| 3db cutoff | **6.96E-03** | 0.02 |
| *Species:Cortical area* | | |
| sag ratio | 8.89E-02 | 0.01 |
| input resistance (SSRn) | 3.13E-01 | 0.00 |
| resonance frequency (fR) | 2.60E-01 | 0.01 |
| 3db cutoff | 3.13E-01 | 0.00 |

Supplemental Table 2 - related to figure 2 and Figure S3

| Variable | Bejamini-Hochberg adjusted p-value | variance explained (η2) |
|---|---|---|
| *Cortical depth* | | |
| sag ratio | **9.26E-07** | 0.12 |
| resting potential (Vm) | **4.20E-03** | 0.04 |
| input resistance (SSRn) | **1.70E-04** | 0.07 |
| resonance frequency (fR) | **7.00E-05** | 0.07 |
| 3db cutoff | **4.20E-03** | 0.04 |
| *Preparation* | | |
| sag ratio | 4.04E-01 | 0.01 |
| resting potential (Vm) | **1.74E-02** | 0.03 |
| input resistance (SSRn) | 9.28E-01 | 0.00 |
| resonance frequency (fR) | 7.99E-01 | 0.00 |
| 3db cutoff | **7.83E-03** | 0.04 |
| *Preparation:Cortical depth* | | |
| sag ratio | 9.98E-01 | 0 |
| resting potential (Vm) | 9.98E-01 | 0.00 |
| input resistance (SSRn) | 9.98E-01 | 0.00 |
| resonance frequency (fR) | 9.98E-01 | 0.00 |
| 3db cutoff | 9.98E-01 | 0.00 |

Supplemental Table 3 - related to Figure 3

| Variable | Bejamini-Hochberg adjusted p-value | variance explained ( η2) |
|---|---|---|
| *Cortical depth* | | |
| sag ratio | **1.00E-06** | 0.13 |
| resting potential (Vm) | 1.72E-01 | 0.01 |
| input resistance (SSRn) | 1.72E-01 | 0.01 |
| resonance frequency (fR) | 3.34E-01 | 0.01 |
| 3db cutoff | **1.92E-02** | 0.04 |
| *Preparation* | | |
| sag ratio | 5.94E-02 | 0.02 |
| resting potential (Vm) | **5.79E-08** | 0.16 |
| input resistance (SSRn) | 5.54E-01 | 0.00 |
| resonance frequency (fR) | 5.54E-01 | 0.00 |
| 3db cutoff | 7.88E-02 | 0.02 |
| *Preparation:Cortical depth* | | |
| sag ratio | 9.17E-01 | 0.00 |
| resting potential (Vm) | 9.17E-01 | 0.01 |
| input resistance (SSRn) | 9.17E-01 | 0.00 |
| resonance frequency (fR) | 9.17E-01 | 0.00 |
| 3db cutoff | 9.17E-01 | 0.00 |

Supplemental Table 4 - related to Fig S5

IT1

| Variable | Bejamini-Hochberg adjusted p-value | variance explained (η2) |
| --- | --- | --- |
| *Cortical depth* | | |
| sag ratio | **1.49E-06** | 0.06 |
| input resistance (SSRn) | **1.61E-09** | 0.15 |
| resonance frequency (fR) | **2.92E-06** | 0.12 |
| *Species* | | |
| sag ratio | **1.87E-28** | 0.44 |
| input resistance (SSRn) | **4.13E-09** | 0.14 |
| resonance frequency (fR) | **1.49E-07** | 0.16 |
| *Cortical depth:Species* | | |
| sag ratio | **3.40E-02** | 0.02 |
| input resistance (SSRn) | 6.23E-01 | 0.00 |
| resonance frequency (fR) | 5.94E-01 | 0.00 |

IT3

| Variable | Bejamini-Hochberg adjusted p-value | variance explained (η2) |
| --- | --- | --- |
| *Cortical depth* | | |
| sag ratio | **1.20E-02** | 0.05 |
| input resistance (SSRn) | **7.49E-03** | 0.07 |
| resonance frequency (fR) | **7.49E-03** | 0.09 |
| *Species* | | |
| sag ratio | **4.37E-03** | 0.08 |
| input resistance (SSRn) | 9.63E-01 | 0.00 |
| resonance frequency (fR) | 6.60E-01 | 0.01 |
| *Cortical depth:Species* | | |
| sag ratio | 7.86E-01 | 0.00 |
| input resistance (SSRn) | 7.86E-01 | 0.00 |
| resonance frequency (fR) | 6.57E-01 | 0.02 |

Supplemental Table 5 - related to Figure 5 and Fig. S7
